# Cell state transition analysis reveals a causal network that controls critical limb ischemia

**DOI:** 10.1101/2024.02.09.579659

**Authors:** Oleksii S. Rukhlenko, Joseph M. McClung, Anna Tuliakova, Brian H. Annex, Boris N. Kholodenko

**Affiliations:** Systems Biology Ireland, School of Medicine, University College Dublin, Dublin, Ireland; Department of Physiology Brody School of Medicine, East Carolina University; Department of Medicine, Medical College of Georgia at Augusta University; Conway Institute of Biomolecular & Biomedical Research, University College Dublin, Ireland; Department of Pharmacology, Yale University School of Medicine, New Haven, USA

## Abstract

Increased endothelial cell (EC) permeability significantly contributes to peripheral arterial disease and its severe manifestation, critical limb ischemia (CLI). CLI remains refractory to current pharmacological interventions, often requiring limb amputation and showing how limited our knowledge is about signaling pathways that control EC states. Here, we utilize a two-pronged approach, employing cell State Transition Assessment and Regulation (cSTAR) analyses to both the published knowledge (e.g., LINCS data portal) and our own transcriptomics data collected from CLI patients and healthy adult (HA) control. We identified CLI and HA transcriptomics patterns and reconstructed causal network that controls transitions between EC states associated with CLI and HA. Contrary to general belief, our study demonstrates that VEGF administration slightly promotes the CLI EC states, indicating VEGF ineffectiveness in enhancing perfusion of CLI tissue.

Based on the unveiled control network, we suggest targets whose inhibition would reverse CLI EC states closer to HA states.

## Introduction

Peripheral arterial disease (PAD) is one of the major complications of systemic atherosclerosis where occlusions in the large arteries reduce the supply blood to the leg(s)^1^. PAD afflicts more than 200 million world-wide and PAD is expected to become an even greater healthcare problem in the years ahead as the major epidemiological drivers for PAD continue to increase in prevalence^1,2^. Patients with PAD suffer from walking limitations and PAD is the number one cause of amputations in adults^3,4^. No therapies are available to improve lower limb perfusion in PAD^3,4^. In patients with PAD the quantity of blood flow distal to the occlusion is dependent on the response of the blood vessels and muscle beyond the occlusion PAD making the assessment of human tissue essential in the study of PAD^5^.

Critical limb ischemia (CLI) represents a severe manifestation of PAD, characterized by significant morbidity and the risk of limb loss, underscoring the existing unmet and urgent need for effective treatments^4,5^. Current CLI treatments involve surgical operations to improve blood supply and pharmacological interventions to prevent thrombosis events. Improperly enhanced vascular permeability is now appreciated to play a significant role in the pathobiology of CLI^6,7^. The increased leakage of blood components adds to inflammation and tissue damage, further impairing limb perfusion and exacerbating ischemic conditions. Existing re-vascularization treatments involving vascular endothelial growth factor (VEGF) have proven to be inefficient. Agents designed to increase blood vessel growth likely increase vascular permeability^8^ and none of the current standard of care treatments specifically targets endothelial cells (ECs) to mitigate vascular permeability.

The present research endeavors to unveil and comprehend the control mechanisms governing EC states in CLI, aiming to identify novel targets for more effective pharmacological strategies. To achieve this, we employed cell State Transition Assessment and Regulation (cSTAR) analysis^9^ to reconstruct and understand causal EC networks that control CLI. We acquired RNA-seq data obtained from 5 patients with chronic limb threatening ischemia (CLTI that is the late CLI stage) and 5 healthy adults (HA). cSTAR analysis of RNA-seq patient’s data merged with publicly available drug perturbation responses of ECs from the LINCS database^10,11^ helped us identify specific EC states in CLI vs. HA. Additionally, it facilitated the quantification of influence of various drug perturbations on EC states. Employing the cSTAR approach allowed us to reconstruct causal network that govern transitions between EC states associated with CLI and HA. Our results suggest targeted perturbations that interconvert EC states, allowing for designer pharmacological approaches to influence cellular pathways and mechanistically underpin pharmacological interventions.

## Results

### Data acquisition

Muscle biopsy samples were collected from all patients and control subjects who granted informed consent (see Methods for details) under protocols approved at East Carolina Medical Center and Augusta University. Muscle biopsies from the lateral head of the gastrocnemius muscle from control subjects were obtained by percutaneous biopsies, using methods previously described^12^. A total of 5 samples from patients with CLTI and 5 from control subjects were acquired. After RNA extraction, RNA sequencing data were acquired as described in Methods.

### Mapping transcriptional states and quantifying CLI and HA phenotypes

In the initial cSTAR step, we applied supervised machine learning, the Support Vector Machine (SVM) algorithm to separate transcriptomic patterns of CLI and HA states. The SVM constructs the maximal margin hyperplane that effectively separates the CLI and HA states. The normal unit vector to this separating hyperplane determines the shortest distance from the CLI states to HA states and is termed the State Transition Vector (STV). Because its direction is selected to indicate a transition from the disease to healthy states, this STV is labeled as STV_CLI-HA_. Principal component analysis (PCA) visualizes projections of the separating hyperplane and the STV_CLI-HA_ into the plane of two first PCA components (Fig. 1).

**Fig. 1.**
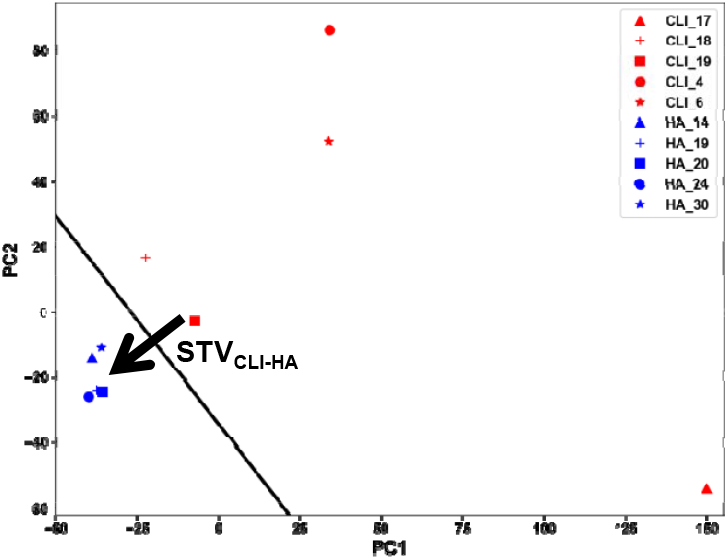
Separation of the transcriptomics patterns of CLI patients and HA controls by SVM and the STV_CLI-HA_ projection into the first two PCA components. Projections of data points of CLI patients (red), the separating hyperplane (black line) and the STV (green arrow) are shown.

Merely using PCA separation between disease and healthy states provides limited insights into the root causes of CLI and potential interventions. Aiming to shift CLI states closer towards healthier states, we exploited a Dynamic Phenotype Descriptor (DPD), which is the distance from a cell state to the CLI-HA separating hyperplane in the transcriptomic dataspace^9,13^. Consequently, for any given patient transcriptome pattern, the DPD_CLI-HA_ score serves as a numerical measure indicating the “healthiness” of the patient’s state. A negative DPD_CLI-HA_ score indicates the presence of CLI, while a positive DPD_CLI-HA_ score, indicates a healthier state.

### Reconstruction of causal network that controls EC states and disease phenotype

To decipher a crucial regulatory network comprising kinases and other protein pathways that govern the transition between CLI and HA states, we exploited perturbation responses of endothelial cells (HUVEC) in the LINCS database^10,11^. An implicit assumption of this analysis is that CLI is driven by the endothelium malfunction and normalizing EC signaling holds the potential to ameliorate the disease state. The adaptation and responses of the endothelium within muscles to injury is critical in PAD^6,12,14^. Though myocytes comprise the greatest cell mass in muscle, it is not unreasonable to conclude that ECs drive the responses of the myocytes and the muscle as the whole in PAD. Generating functional, stable blood vessels has to be a goal of human therapeutics and generating tumor-like vessels within an ischemic muscle, whether in a mouse or a human, would be expected to be detrimental^15,16^.

Leveraging the LINCS HUVEC datasets^10,11^, we established transcriptomic signatures of drug targets that include kinases and other proteins (see Methods section and ref^17^). Based on the data, we selected components of a core network that are associated with the control of transitions between CLI and HA states. Because the DPD_CLI-HA_ score describes the phenotype as a dynamic summary of the cell-wide network (proteins, mRNAs, etc), we included the DPD_CLI-HA_ as a node in the core network. This enables us to systematically investigate the impact of all core network pathways, both individually and in combination, on the transitions between CLI and HA states. The selection of signaling modules is predominantly data-driven, but involves a partially manual process, allowing the flexibility to add or remove modules as necessary^9,13^.

Subsequently, we merged the STV_CLI-HA_ derived from the patient data and the LINCS perturbation data to compute both the changes in the DPD_CLI-HA_ and the variations in the activities of core network components, utilizing their transcriptomic signatures (see Methods). Both these alterations served as the global response input to a cSTAR network inference algorithm, which is based on the Bayesian formulation (BMRA) of Modular Response Analysis (MRA)^9,18^. Then, BMRA inferred the directions and strengths of causal connections, including feedback loops, between network pathways and direct connections from each of these pathways to the DPD_CLI-HA_ module. The causal connections quantify the direct, local influence of each network module on other modules, rather than capturing the global responses of these modules. Global responses manifest through correlations or associations that arise as a result of the propagation of direct responses throughout the network.

Fig. 2 illustrates reconstructed causal connections of the core network that controls EC state transitions. As observed, VEGFR activates ERK and PI3K that in turn activates PAK. PKC activates VEGFR and PI3K and inhibits the Aurora kinases A and B (AURCA/B), which are also inhibited by CDK1/2 that is upstream of AURCA/B during cell cycle progression. AURCA/B appeared to be an immediate CLI driver, because of its negative direct effect on the DPD_CLI-HA_ module, hampering a transition from CLI to HA. Other immediate drivers of CLI include HDAC and PAK, which both have negative direct effects on the DPD_CLI-HA_. Notably, a positive feedback loop between AURCA/B and PAK reinforces driving CLI states by these kinases. Immediate drivers of HA include PKC, CDK1/2, PI3K and TGFbR, which have positive influence on the DPD_CLI-HA_, facilitating the CLI – HA transition. Notably, because of multiple feedback loops and crosstalk between signaling pathways, their immediate direct effect on CLI – HA transition can be superseded by their global influence that stems from network-wide effects.

**Fig. 2.**
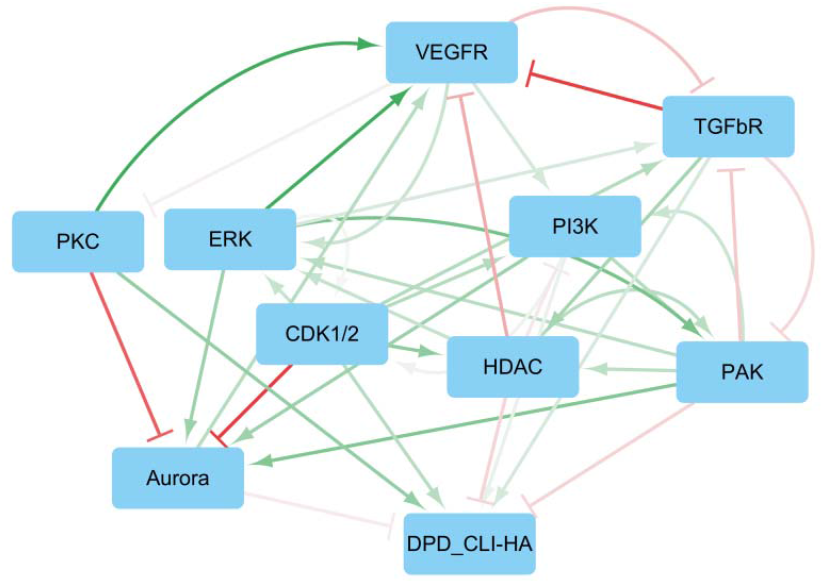
Causal network controlling a transition between CLI and HA states associated with ECs. Green lines with arrowheads indicate activation, red lines with blunt ends show inhibition, the line widths correspond to the absolute values of the interaction strengths.

### Suggesting potential drug targets to ameliorate CLI

MRA also enables us to calculate the global effects of each network module on other modules, including the DPD_CLI-HA_, which are termed global impacts. Fig. 3 displays a bar plot illustrating the global impacts of each pathway of the controlling network on the DPD_CLI-HA_. Similarly as for direct impacts, the negative value indicates that a pathway overall drives the CLI state, whereas positive value indicates that a pathway drives the HA state. Surprisingly, VEGFR weakly drives CLI state, albeit not directly. Instead, it drives CLI state by activating downstream targets, ultimately leading to the activation of PAK. The uncovered VEGF contribution to CLI-HA transition supports clinical evidence that VEGF therapies do not help patients with PAD^16^.

**Fig. 3.**
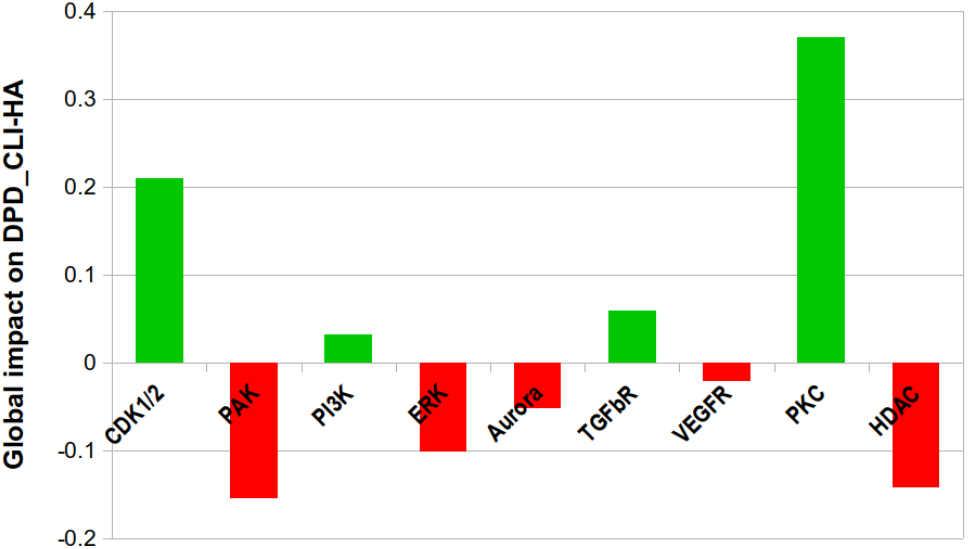
Network-wide, global impacts of signaling pathways on the transition from CLI state to HA state. Green and red columns depict contributions of pathways that facilitate or hamper transitions from CLI state to HA state.

Afterwards, we used transcriptomics signatures of pathways in the controlling network to quantify the levels of their signaling activities in CLI patients and HA controls. The results are presented at the Fig. 4. Surprisingly, although the Aurora kinase serves as one of the drivers of the CLI state, the activity of this kinase in CLI patients is significantly lower compared to HA patients. Potentially, suppression of the Aurora kinase could act as a compensatory mechanism when an organism in the CLI state attempts to revert to the HA state. Consequently, we posit that cSTAR has enabled the differentiation of compensatory mechanisms from overall phenotypic changes. Given the substantial suppression of Aurora kinase activity in CLI patients, our analysis indicates alternative potential drug targets for CLI, including PAK and HDAC (Figs. 2 and 3). Summarizing, the cSTAR approach has allowed us to identify the drivers of CLI states and potential novel drug targets, independent of compensatory effects, thereby enhancing our understanding of disease progression.

**Fig. 4.**
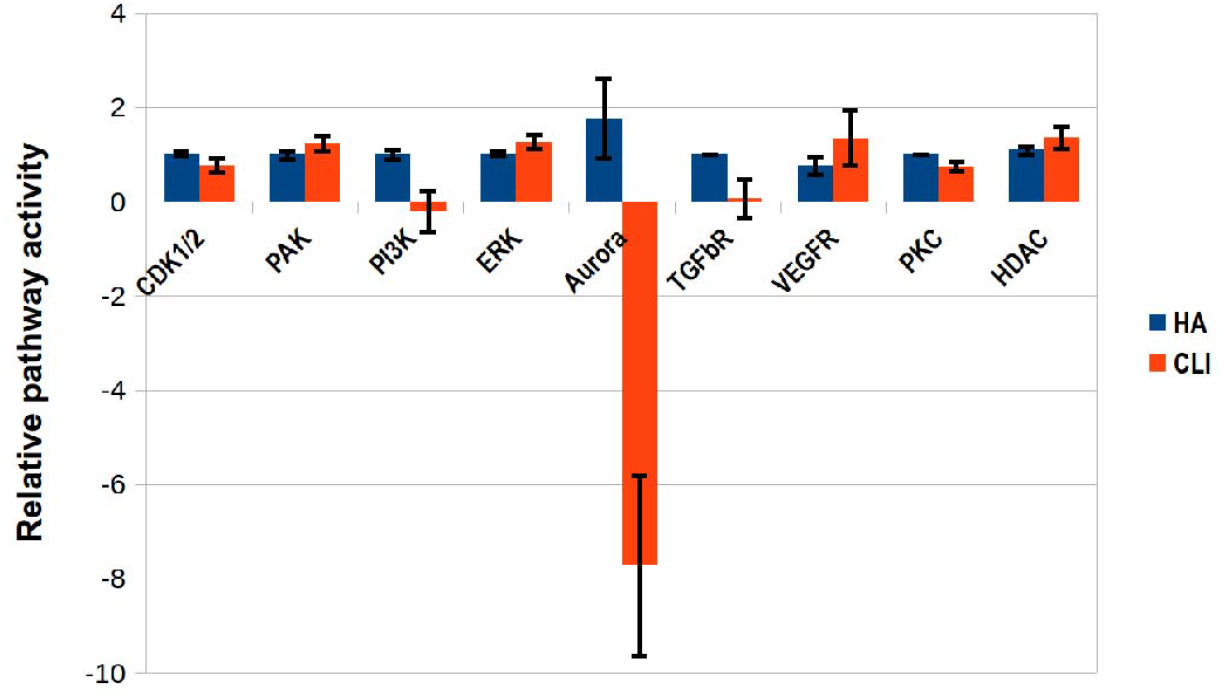
Relative pathway activities pathway in CLI and HA states. Red and blue indicate pathway activities in CLI and HA states, respectively.

## Discussion

The analysis of human tissue is essential for advancing our understanding of human disease. Rodent models of PAD are invaluable but the complexity of the human condition is such that models can be used to address targeted questions but the disease in its entirety. Given the importance of the endothelium to PAD, bulk whole-muscle RNA-seq data was obtained and submitted for cSTAR analysis designed to elucidate EC transcriptomic profiles. Standard analysis of mRNA-seq from muscle from patients from ECU has been previously reported and “standard” analyses of patients with CLTI vs. controls show striking differences^12^. Yet, the uncovered differences lack actionable insights into the critical pathways governing the CLTI patient states compared to HA states. A main pillar of cSTAR is to analyze omics data to map cell states, reveal molecular features that govern their transitions, and comprehend the signaling networks controlling them. Reconstructing and understanding control networks necessitate systemic perturbations that patient data alone cannot provide. Such data can only be obtained ex-vivo using EC lines or muscle organoids. Here we employed a rich dataset acquired by the Broad Institute on transcriptomic responses of ECs (HUVEC cells) to a variety of drug perturbations^10,11^, integrating it with our transcriptomic data collected from patients. The patients’ data allowed us to quantify the states ECs in patients based on a healthiness metric, using the DPD_CLI-HA_ score. Combining this with the in vitro perturbation data of HUVEC, we were able to infer the directions and strengths of causal connections within the control network.

Further cSTAR analysis unveiled potential drug targets whose inhibition could potentially ameliorate CLTI conditions by enhancing perfusion.

All reported conclusions from these studies need to be validated in larger samples sizes and with validation of specific measures within ECs in muscle tissue that are predicted alterations from the analysis. What was perhaps most striking of the results of this cSTAR analysis, was that data from the CLTI vs. controls had similarity to proliferating vs. non-proliferating ECs; or ECs subjected to VEGF stimuli. Increasing muscle perfusion is the goal of treatment for PAD. Merely growing blood vessels is not a goal of therapy. At an extreme, growing tumor-like blood vessels in ischemic muscle is unlikely to be beneficial to patients with PAD and could easily be viewed as harmful. cSTAR analysis has effectively captured this pivotal point and suggested potential targets to normalize the state of endothelium.

## Methods

### Data acquisition

Patients were recruited from the vascular surgery clinics at East Carolina University (ECU) Medical Center in Greenville, NC. Muscle biopsy samples were obtained on all patients and controls who provided informed consent under approved protocols at ECU and de-identified materials were sent to Dr. Annex’s laboratory at the University of Virginia (UVA) and Medical College of Georgia (MCG) at Augusta University where RNA-sequencing was performed and analyzed.

In total samples from 5 patients with chronic limb threatening ischemia (CLTI) and 5 controls were studied. The 5 controls were age 62 – 69 years, 3 men and 2 women, 2 were active smokers. The CLTI patients were age 55 – 76 years, 2 men and 3 women, none were active smokers and 2 had diabetes both with normal hemoglobin A1cs.

Muscle biopsies from the lateral head of the gastrocnemius muscle from control subjects were obtained by percutaneous biopsies, using methods previously described^12^. Muscle biopsies from patients with CLTI occurred in the operating room and were obtained the time of amputation that was performed without the use of a tourniquet. Samples included only ones were where tissue oxygenation was sufficient to allow wound healing in PAD patients. Muscle biopsies were collected form the same anatomical location of the gastrocnemius muscle as the percutaneous muscle biopsies from HA and IC patients. An obvious concern is that muscle from patients with CLI will be badly degraded. Muscle viability and structural/morphological integrity was verified to exclude necrosis and samples were assayed for mitochondria (citrate synthase/soluble protein and mitochondrial:nuclear DNA) activity. This method has previously produced viable biological sampling in PAD patients within the McClung lab^12^.

At the University of Virginia, RNA-seq libraries were constructed from 500 ng of total RNA using the TruSeq 2 Stranded Total RNA prep kit (Illumina, San Diego, CA). The resulting barcoded libraries were sequenced on an Illumina NextSeq 500. Quality control of the resulting sequence data was performed using FastQC^19^. Raw reads were aligned against hg38 human reference genome using Rsubread R package. The resulting raw counts were used to calculate differentially expressed genes using DeSeq2 R package. RNA sequencing data will be deposited following a journal submission.

### Building the STV and using the LINCS data

Using a cSTAR approach, we separated of CLI and HA states and determined the STV_CLI-HA_ that indicates the direction of the shortest path from the CLI states to HA states in the transcriptomic dataspace, as described previously^9,13,17^. Further, the STV_CLI-HA_ is designated by. Then, we utilized LINCS perturbation data to infer causal drivers of CLI and HA states. The LINCS database contains a large number of HUVEC transcriptomic responses to drug perturbations, allowing to perform inference of causal connections between the drug targets.

To make the STV transferrable between different datasets, the STVs is built in the space of log-fold changes (LFCs). For patient data, we used the average of all HA states as a reference. In LINCS data, the reference state used for calculating the LFCs is a static condition for HUVEC cells without any perturbations. To estimate phenotypic changes brought about by drug perturbations, we calculated the dot products of the STV_CLI-HA_ and the vectors of log fold changes obtained from the LINCS data.

Namely, we calculate dot products of the LFC vector (***x***_*j*_), which is determined from the LINCS data, and the STV_CLI-HA_ (***n*)**, to find the element of the global response matrix (*R*_*DPD,j*_) for the j-th_CLI-HA,_ perturbation corresponding to the DPD_CLI-HA_ module, as follows,

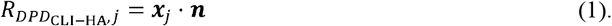

### Global responses of core network modules

If a direct measurement of the core network module activity can be done, the LFC in the expression of the gene *i* in response to the *j*-th perturbation are used for *R*_*ij*_ value. When direct measurements of the activity of the i-th node are unavailable, such as in transcriptomic assessments of responses to enzyme inhibitor treatments, we apply the following procedure. A relative activity of the of the i-th module after j-th perturbation (*f*_*ij*_) is calculated as a linear function of the transcriptomic LFCs ( *x*_*kj*_) as follows,

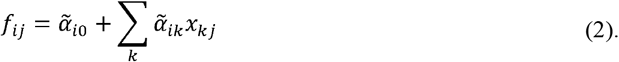

Coefficients 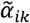,which are used to infer the relative activity of the i-th module, are found using linear regression over the function *g*, which describes responses of the correspodong enzyme activity to an inhibitor. In the simplest case of a monomeric enzyme this function is expressed as follows^17^,

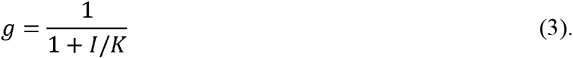

Here is the inhibitor concentration and *K* is the IC50 dose for this inhibitor, taken from the literature data. If multiple inhibitors of the same target are applied, the coefficients 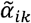 are found by regressing the function *g* over all doses of all inhibitors applied. The coefficient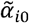 is set to 1 for all modules, because for the control (no inhibitors) data points, all LFC, *x*_*kj*_, are close to zero, while the value of the function *g* is equal to 1.

To satisfy the MRA modular insulation condition^20^, vectors of the coefficients 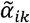 (excluding the intercept coefficient 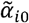) for different modules must be mutually orthogonal. Additionally, we have penalized the negative pathways activities and added the Lasso regularization to minimize the number of potential solutions to ideally obtain a unique solution. In brief, we infer the coefficients 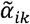 by finding the global minimum of the following function,

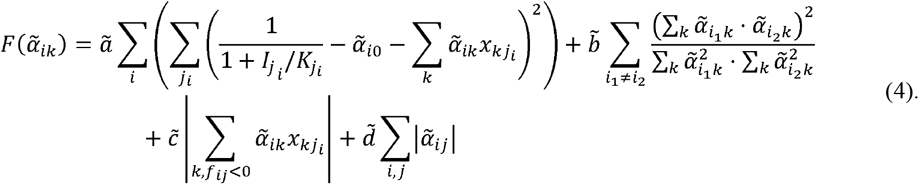

Here the first term describes the linear regression over enzyme activity for each network module, the second term introduces the orthogonality condition, the third term penalizes the negative pathway activities and the fourth term introduces the Lasso regularization. The hyper-parameters, 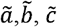 and 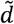, provide a means to balance the priority among different terms when it is not possible to satisfy all conditions simultaneously.

Taking into account that 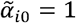, Eq. 4 will read,

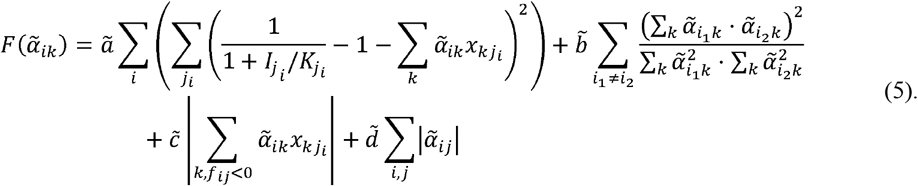

When coefficients 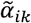 are determined, the elements of global response matrix *R*_*ij*_ for core network modules can be calculated from Eq. 2 as follows,

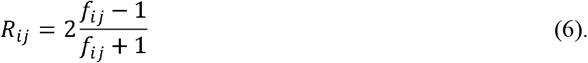

Here we imply that for the control data point *f*_*ij*_ =1

It is important to highlight that while the STV (***n***) is normalized to have a unit length, the vectors 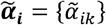 do not have unit lengths and must not be normalized, as doing so would violate the regression conditions. Completing the entries, *R*_*ij*_, for both the core network modules and the DPD module(s) marks the completion of the transcriptomic data preparation for the BMRA inference in cSTAR.

### Relative pathway activities of different patients

In the analysis of patient data, we employed the log-fold changes in gene expression relative to the averaged HA states as inputs for Eq. 2. This allowed us to calculate the relative activities of core network modules for both HA and CLI groups. It is essential to emphasize that activities calculated in this manner only hold significance as relative values, and they estimate the relative activity of each signaling module within a patient compared to another patient.

